# Visualization of spatial distribution of hemoglobin with various oxygen saturations in small animals using a photoacoustic imaging scanner with a hemispherical detector array

**DOI:** 10.1101/2023.06.19.545650

**Authors:** Yasufumi Asao, Ryuichiro Hirano, Kenichi Nagae, Hiroyuki Sekiguchi, Sadakazu Aiso, Shigeaki Watanabe, Marika Sato, Takayuki Yagi, Shinae Kizaka-Kondoh

## Abstract

**Significance:** Photoacoustic (PA) imaging has garnered considerable attention due to its capability to render vascular images in a label-free manner. Specifically, devices employing a hemispherical detector array (HDA) have been heralded for various clinical applications, owing to their potential to yield high reproducibility three-dimensional images. While high-resolution models utilizing high-frequency sensors have been introduced for animal experimentation, their evaluation has been constrained to a single wavelength. In this study, we demonstrate the applicability of in vivo mouse models for visualizing body oxygen saturation distribution using dual wavelengths.

**Aim:** With the aid of our uniquely developed device and analysis software, our primary objective is to map the spatial distribution of the hemoglobin oxygen saturation coefficient (S-factor) through non-invasive in vivo imaging. Subsequently, we aim to observe the temporal alterations within this distribution, specifically assessing changes in hemoglobin oxygen saturation in both normal and tumor vessels over time.

**Approach:** High-quality S-factor images were obtained by integrating a newly developed scanning sequence for high contrast with alternate two-wavelength irradiation. Following validation with phantoms, in vivo images were procured in mice. Sequential scanning of the same mouse yielded information about temporal changes. S-factor evaluation was conducted with our photoacoustic image viewer to analyze trends in hemoglobin oxygen saturation.

**Results:** High-contrast images were achieved by increasing the number of integrations during scanning. S-factor images were acquired using both healthy and tumor-bearing mice. Vessels within the liver and kidneys were distinctly reconstructed, and differences in oxygen saturation discriminated between arteries and veins. Repeated measurements on the same mice, both live and post-euthanasia, provided spatiotemporal information, such as a decrease in oxygen saturation after euthanasia or a precipitous drop in oxygen saturation inside the tumor nine days post-cell line transplantation.

**Conclusions:** By analyzing S-factor images using a photoacoustic imaging system designed for animal experiments, we succeeded in discerning variations in in vivo oxygen saturation. The custom-built system holds promise as a versatile tool for diverse basic research endeavors, as it can seamlessly interface with human clinical applications.

## 1 Introduction

Assessing oxygen saturation is pivotal in evaluating disease pathogenesis and progression. Within solid tumors, hypoxic conditions are known to correlate with prognosis^1^; thus, determining the level of oxygen saturation in tumors is crucial. Solid tumors typically present hypoxic environments due to the rapid proliferation of cancer cells coupled with abnormal angiogenesis^2^. Tumor hypoxia plays a role in several malign mechanisms of cancer, including radioresistance, drug resistance, immunosuppression, and metastasis, contributing to poorer patient prognosis^3-6^. As such, intratumoral hypoxia stands as an important prognostic factor^7^ and is instrumental in selecting appropriate treatment. Immunohistochemical analysis, detecting hypoxia-inducible factor-1α (HIF-1α) protein and pimonidazole-binding macromolecules as hypoxia markers, is employed to evaluate tumor hypoxia^8,9^. Alternatively, tumor hypoxia can be gauged by measuring hemoglobin oxygen saturation in the blood. For instance, assessing hemoglobin oxygen saturation in primary breast cancer using diffuse light spectroscopic imaging has demonstrated that oxygen saturation correlates with the pathologic response to neoadjuvant chemotherapy^10^. Therefore, determining oxygen saturation within the tumor is vital for predicting treatment efficacy.

Photoacoustic (PA) imaging is an emerging hybrid technique that merges the high contrast and spectral unmixing capabilities of optical imaging with the high spatial resolution of ultrasound, making it an active area of research due to its potential to provide label-free vascular images^11-13^. One approach to enhance the quality of PA images involves designing the sensor to encompass the imaging target, thereby mitigating the limited view issue^14^. Our team has pioneered a high spatial resolution device that employs a hemispherical detector array (HDA) to perform scans in the horizontal plane^15-18^. We are currently conducting clinical research on human subjects using this device^19-21^. The device is now approved by the Pharmaceuticals and Medical Devices Agency (PMDA) in Japan for use as a medical device.

We designed a photoacoustic device similar to the one used for human subjects, employing the same HDA configuration, and reported on its application to animals^17^. The wavelength evaluated in that study was a single wavelength of 797 nm. In this study, utilizing our newly developed photoacoustic device for animal experiments, we successfully obtained an image correlated with oxygen saturation (S-factor image) from images captured at two wavelengths. This system enabled us to procure S-factor images of living organs and tumors in mice transplanted with tumor cells, and we present our findings in this paper.

## 2 Materials and methods

### 2.1 Device configuration

The main configuration of the experimental apparatus for animals developed in this study is akin to the bed-type device used in our past human clinical studies. It consists of a laser irradiation unit, a hemispherical sensor to receive photoacoustic waves, and a data processing unit. In a previous study^17^, we utilized lasers capable of emitting wavelengths of 797 nm and 835 nm to generate photoacoustic signals but only evaluated the single wavelength of 797 nm. The wavelength of 797 nm is where the optical absorption coefficient of oxygenated and deoxygenated hemoglobin is the same, allowing for the evaluation of vascular morphology.

The apparatus used in this study allows for the choice of irradiation conditions, either repeatedly pulsing the same wavelength of 756 nm or 797 nm, or alternately pulsing between these two wavelengths^16,19^. The repetition frequency was 30 Hz for a single wavelength and 15 Hz for each wavelength when alternating. The measured laser energy per pulse was 16 mJ at 756 nm and 20 mJ at 797 nm. The specifications of these lasers align with those reported in our previous study, with the added feature of being able to utilize a 756 nm wavelength, which is particularly advantageous for visualizing deoxygenated hemoglobin.

The laser is emitted through a lens at the bottom of the hemispherical detector array (HDA), and the laser energy is below the maximum permissible exposure (MPE) level, defined in the laser safety standard, so it is safe for human skin and is not expected to harm animals. The device is equipped with a 512-channel, 5.28 MHz center frequency sensor. The specifications of the sensor are the same as those described in the previous papers.

The PA imaging system used in previous studies^18-20^ was a bed-type system (Fig. S1(a)). In contrast, the system developed in this study is a two-story system with the bed-type device halved and the scanning section stacked on top (Fig. S1(b)). As previously reported, the bed type is suitable for measuring humans in a lying position and offers stable measurement with minimal body movement. However, small animals like mice do not require any space beyond the imaging tray, so image quality remains unaffected. This reduction in footprint to a compact 1.6 m (W) x 1.1 m (D) x 1.1 m (H) makes this configuration ideal for small laboratories. Despite the device’s height being approximately double that of the bed type, it is designed so that an experimenter of 1.7 m height can comfortably conduct animal experiments under his/her chest (Fig. S2).

The device is also equipped with a light-shielding box covering the entire tray, including the laser output section, and a safety mechanism that prevents the laser from emitting light unless the light-shielding cover is closed (Fig. S3). This categorizes it as a Class 1 product, usable safely without the need for protective goggles. The light-shielding box illustrated above can be used to insert a tube for administering inhalation anesthesia to mice during experiments. The tube is inserted from the back of the light-shielding box, and light leakage from the tube insertion port (not shown) is negligibly small.

The tray of the PA imaging system is designed so that acoustic matching water can be filled to a depth of no more than 15 mm, ensuring a good acoustic match with the specimen being imaged. The mouse or other specimen is placed in the tray and submerged underwater. The mouthpiece for inhalation anesthesia is positioned above the water surface using a spacer so that the mouthpiece does not get submerged, and the specimen can continue to breathe (Fig. S4). If necessary, a transparent ultrasound gel can be used instead of water to maintain good acoustic matching with the specimen being imaged.

The device includes a tray filled with water and an HDA, with its sensor surface immersed in the water. These components are separated by a 23-micron-thick polyethylene terephthalate (PET) film, similar to our previous devices for human subjects. A heater warms the water in the device, maintaining a temperature of 37-38°C to mitigate the risk of hypothermia-induced death.

### 2.2 Data acquisition

The unit is fitted with an HDA capable of scanning a large horizontal area. In both modes, the HDA is scanned at a pitch smaller than the image area reconstructed by a single laser shot. Therefore, when focusing on a single voxel, multiple laser shots are performed. This integration of images leads to noise reduction. The number of integrations is 29 in standard (Std.) mode and 470 in high-quality (HQ) mode. Thus, the number of integrations in HQ mode is 16 times greater. It takes 16.5 seconds to capture a mouse-sized image (4 x 6 cm in size) in Std. mode and approximately 210 seconds in HQ mode, which is 12.7 times longer.

The main unit contains a Linux PC equipped with a GPU. The PA signals received by the HDA are transmitted to the PC via the Data Acquisition System (DAS) in the device. They are processed in real-time to generate a three-dimensional (3D) image. The generated 3D image is output to the operation monitor as a maximum intensity projection (MIP) image. Additionally, a built-in digital camera is installed to capture still images at each position during the scan, ultimately creating a composite photograph. This composite photograph can be used to compare the visible light appearance of the mouse with the photoacoustic image generated by the device.

### 2.3 Hemoglobin oxygen saturation approximation

The approximation calculation employs an equation, with absorption coefficients and molar extinction coefficients, to evaluate the oxygen saturation in hemoglobin. However, light propagation within a living body is complex, making it challenging to accurately determine the light fluence. Thus, we use an approximation of the original equation, replacing certain factors with the pixel value of the measured PA image.

In general, the spatial distribution of hemoglobin oxygen saturation is expressed by Equation [1].

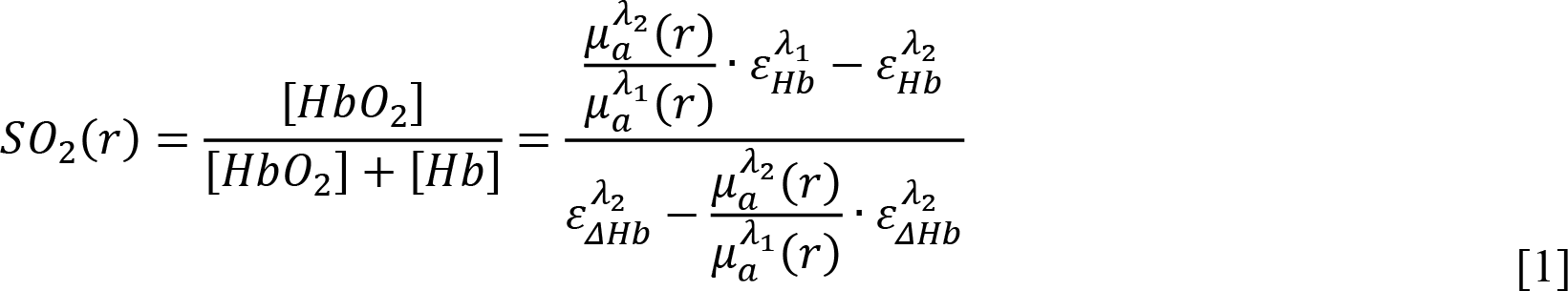

where *μ*_*a*_ is the absorption coefficient and *ε*_*Hb*_ is the molar extinction coefficient of deoxy-Hb and *ε*_Δ*Hb*_ is the difference in the molar extinction coefficients between deoxy-Hb and oxy-Hb (HbO_2_). The subscript *λ*_*i*_ represents the light wavelength and *r* represents the spatial coordinates. From the general formula for photoacoustic signals, *μ*_*a*_ is calculated as 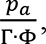 where Γ is the Grüneisen coefficient and Φ is the light fluence reaching the absorber.

However, light propagation in a living body is complex, and it is difficult to accurately determine the light fluence. In other words, *μ*_*a*_ cannot be accurately obtained.

S-factor is an approximation of Equation [1], where *μ*_*a*_ is replaced by the pixel value *p*_*a*_ of the measured PA image and changed to Equation [2].

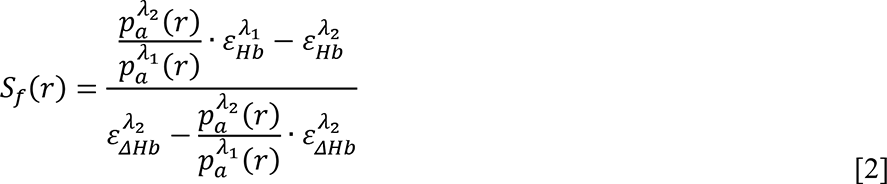

In a previous report^18^, the initial sound pressure distribution at each wavelength was calculated and imaged assuming that the light intensity ratio at each wavelength was the same. This time, with the software update, images are automatically obtained after being corrected by the average irradiation light intensity of each wavelength during the scan. This allows for compensation of the difference between the two wavelengths of energy emitted from the laser, thereby aligning it closer to the true value. Nonetheless, it should be noted that the absolute value is not guaranteed to be correct because the optical fluence in vivo is not evaluated.

### 2.4 Image viewing and analysis

The images generated by this device can be loaded into a bespoke Windows application we developed, called the PAT Viewer. The PAT Viewer is the successor software to a previously used application, KURUMI^22^, demonstrating improved capabilities in presenting voxel data more accurately. With this novel tool, the images generated by our system can be used for various types of analysis and processing. For instance, the PAT Viewer can automatically trace a specific vessel when indicated, or manually designate an arbitrary block region and compute the average value of the S-factor within that defined range. When quantifying the S-factor, a weighted average value, factored by the photoacoustic image intensity at 797 nm, is utilized.

### 2.5 Imaging target

#### 2.5.1 Chart phantom

For testing the instrument’s still image mode, we used two phantoms: a USAF1951 resolution chart and an ISO12233 chart23. As the former has been previously reported^17^, we report on the latter in this report. The ISO12233 chart used in this experiment was printed on a 3 mm thick A4-sized acrylic plate by a dedicated inkjet printer (MIMAKI UJF-6042MkII, Japan) and contains the patterns specified in the standard (Fig. S4). This phantom was used to evaluate the performance of the device in still image mode.

#### 2.5.2 Healthy mouse

The system was tested on white (or albino) hairy mice of the B6 and BALB/c strains (Charles River Laboratories Japan, Japan). Experiments on healthy mice were performed using 9-week-old female mice. Just before imaging, a hair removal cream (Veet Botanicals Hair Removal Cream for Sensitive Skin, Reckitt Benckiser, UK) was applied to the body surface of the mice to completely remove hair around the imaging area.

#### 2.5.3 Orthotopic breast cancer mouse model

The 4T1 murine breast cancer cell line was purchased from ATCC (Manassas, VA). 4T1/Fluc cells were established as previously reported^24^. Cells were maintained with 5% fetal bovine serum (FBS)-Dulbecco’s Modified Eagle’s Medium (Nacalai Tesque, Kyoto, Japan) supplemented with penicillin (100 units/ml) and streptomycin (100 mg/ml), cultured in a 5% CO_2_ incubator at 37 °C, and regularly checked for mycoplasma contamination using a mycoplasma detection kit (Lonza, Basel, Switzerland). All cell lines were independently stored and recovered from the original stock for each experiment.

Cell suspensions of 4T1/Fluc (3.0 × 10^5^ cells) in PBS were mixed with an equal volume of Geltrex^®^ (Invitrogen, Waltham, MA) and injected into the fourth mammary gland fat pad of 8–9-week-old BALB/c albino female mice.

Tumor size was measured manually by visual inspection of the long and short diameters and the location of the tumor center. Hair removal was performed immediately before imaging as in the case of healthy mice.

### 2.6 Scanning method for mice

Using the MK-AT210D anesthesia machine for small animals (Muromachi Machinery, Japan), a mouse was allowed to inhale a gas mixture of 2% isoflurane (Fujifilm Wako Pure Chemical Corporation, Japan) and air. During scanning, water-absorbent sponges approximately 1 cm thick were placed on the legs and tail to suppress body movements, irrespective of the mouse’s posture. If the mouse was supine, a sponge was placed on the torso to suppress body movements of the mouse. Scanning was performed one mouse at a time. The cervical dislocation method was used to euthanize the mice.

### 2.7 Ethics

Animal experiments in this study were conducted with the approval of the Tokyo Institute of Technology’s Ethics Committee for Animal Experiments (#D2021018).

## 3 Results

### 3.1 ISO12233 chart phantom

Figure 1(a) presents a Maximum Intensity Projection (MIP) image, which is an overall representation of the ISO12233 chart taken in the standard (Std.) mode with our system. The chart was completely captured without any distortion. The yellow rectangular area in the center of the image in Fig. 1(a) was the focal point of the evaluation, and the chart pattern was scanned using two modes: Std. and High Quality (HQ) modes. The area of 40mm x 60mm, which includes the yellow rectangle, was scanned, and a specific area of 25mm x 50mm was cropped from the acquired image. Figure 1(b) displays a magnified MIP image of the yellow area in Fig. 1(a), comparing the differences in outcomes due to the distinct scanning modes. For the background area lacking a chart pattern, our attention was on the Std. mode (A) and the HQ mode (B). The black level in HQ mode (B) is darker than in Std. mode (A), suggesting improved contrast. Figure 1(c) illustrates all the pixel values from the background areas marked by (A) and (B) in Fig. 1(b) extracted and graphed as a boxplot. This graph enables a numerical comparison of each noise level. We performed a Welch’s t-test to numerically compare the scanning modes. The p-value was significantly smaller than 0.01, indicating that the HQ mode yields considerably lower pixel values. Comparing the average pixel value of the background area in both modes, the HQ mode was found to have a value 4.6 times smaller than that of the Std. mode. Furthermore, the pixel values of the signal areas where the pattern was printed were about 60,000 in both cases, indicating that the outcomes differ by a contrast factor of 4.6.

**Fig. 1.**
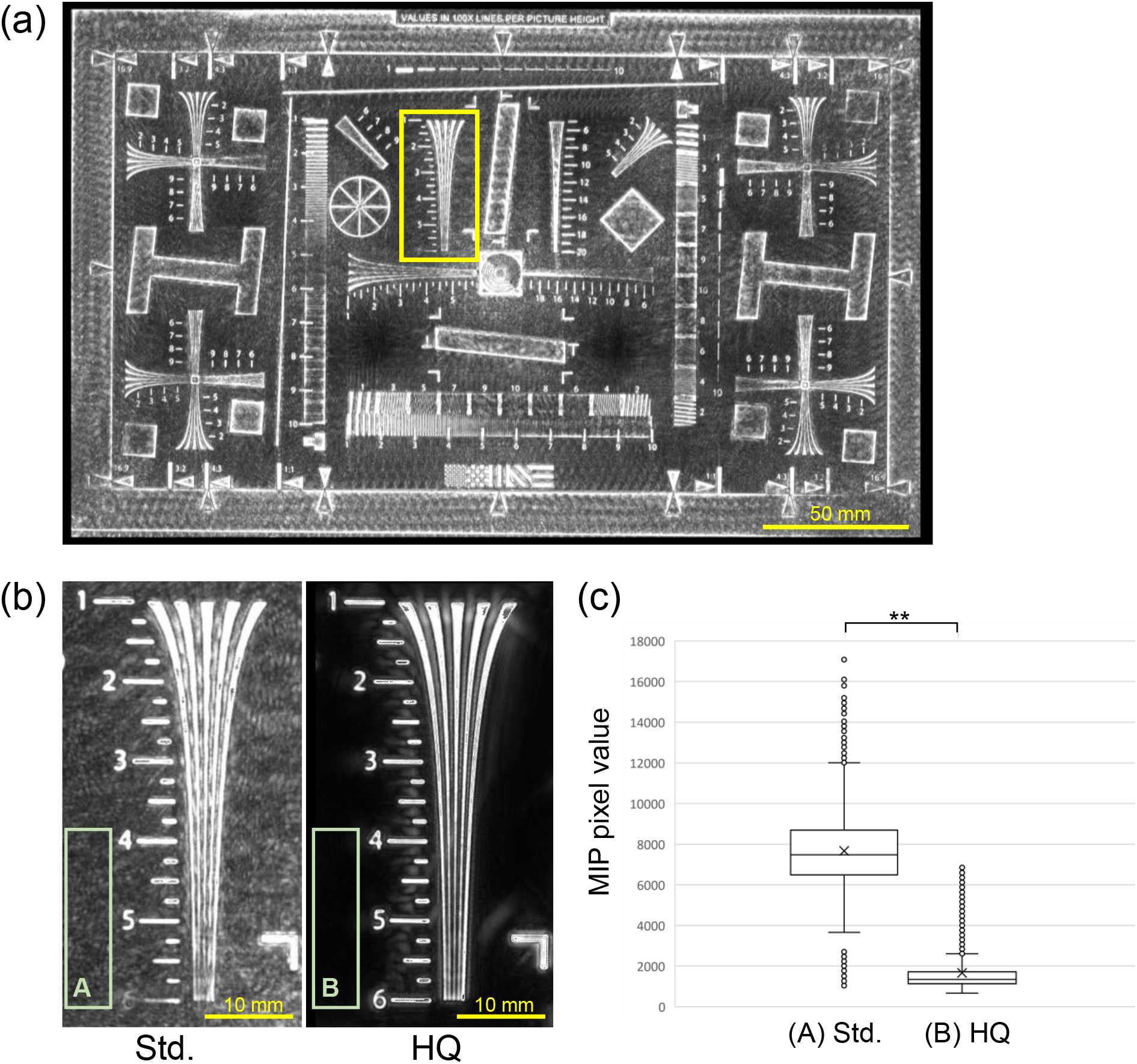
Photoacoustic images acquired using the ISO12233 chart phantom and the resultant numerical analysis. (a) Maximum intensity projection (MIP) image of the entire chart scanned in Std. mode, demonstrating that the full A4-sized chart was imaged without distortion. The yellow rectangle is analyzed in (b) and (c). (b) MIP images scanned in Std. mode (left) and HQ mode (right). The pixel values within the high-intensity region, where the chart pattern exists, are nearly identical in both modes. The area marked by the green rectangle in the lower left (A, B), where there is no chart pattern, shows that the HQ mode appears darker than the Std. mode.. (c) Values of MIP imagewith green squares for Std. mode (A) and HQ mode (B) in (b) were analyzed and shown in boxplots. n=10000. **p<0.01

### 3.2 Photoacoustic imaging of healthy mice

We assessed mice by scanning them using the HQ mode, which offered a high signal-to-noise ratio in phantom experiments. The hair in the measurement area was removed using a hair remover shortly before scanning. BALB/c mice were scanned in a prone position. The imaging results at a wavelength of 797 nm are shown in Fig. S6 and Fig. 2. Figure S6 exhibits the PA and digital camera images of the entire mouse body captured with the current device. Figure 2 depicts a full-body image scanned from the abdomen side and an image specifically focusing on the liver. Figure 2(b) shows a magnified PA image of the area enclosed by the red square in Fig. 2(a), which is a full-body PA image. Based on anatomical observations such as in vivo location and vascular geometry, the organ displayed in Fig. 2(b) was identified as the liver. Figure 2(c) is a microscopic image of the liver removed from the same mouse, dissected after photoacoustic imaging. When compared to the anatomical photograph of the vascular structure of the lower end of the liver displayed in Fig. 2(c) at the same scale, a similar vascular network was found between the PA image and the anatomical photograph. The estimated locations of similar vascular branches were also measured on a microscale, and it was hypothesized that vessels with a diameter of at least 100 microns or less could be reconstructed. Figure 2(d) is a colorized image of the full-body depth information. This uses the grey-scale image in Fig. 2(a) as intensity information and assigns depth (z-axis) length as color information. It should be noted that this is colored with the horizontal plane as the xy-plane and the bottom edge of the image as z=0, and does not necessarily represent the depth length of the mouse organism. For the in vivo depth-length assessment, the area enclosed by the yellow square in the middle of Fig. 2(d), i.e., near the mouse liver, was selected. Figure 2(e) is an MIP image of a rectangle with a 5 mm x 26 mm yellow rectangle near the liver in Fig. 2(d), trimmed in three dimensions and viewed from the cross-sectional direction (i.e., caudad). The yellow triangles mark the positions of the deepest and shallowest presumed areas in the image, which are considered to be the biological signal different from the noise. The spotted image displayed by the triangles near the skin surface was observed, and a linear structure was observed in the deep area, which was presumed to be a blood vessel, suggesting that a vascular structure with a depth of approximately 10 mm was reconstructed (Fig. 2(e)).

**Fig. 2.**
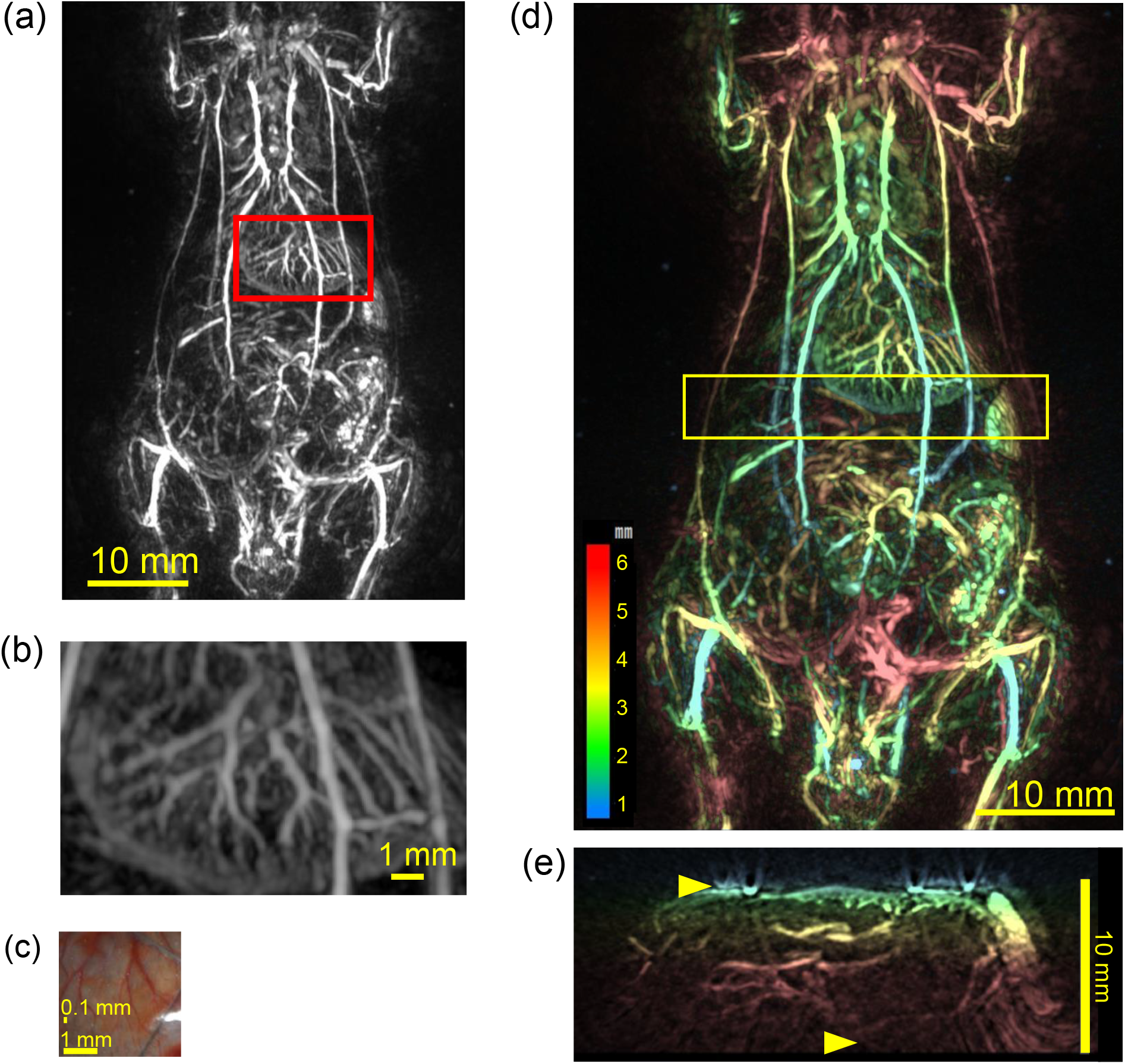
Photoacoustic image of a mouse scanned from the ventral side at a wavelength of 797 nm. (a) MIP image of a photoacoustic image of a mouse scanned in a 4 cm × 6 cm area. The central red rectangle indicates the liver’s location. (b) Enlarged image of the area enclosed by the red square in (a). (c) Micrograph of a liver excised from the mouse in (a). (d) Depth-colored view of (a). Blue indicates the shallow region near the laser emission area and HDA, while red signifies the deeper region. The yellow rectangle in the figure indicates the area of tomogram analysis in (e). (Video-1, MP4, 9.3MB). (e) MIP image of the tomogram viewed from the caudal side. The two yellow triangles represent shallow and deep vascular-like signals, respectively.

Next, Fig. 3(a) presents S-factor images, measured and calculated at two wavelengths, 756 nm and 797 nm, in addition to the black and white imaging at 797 nm depicted in Fig. 2. The vascular network within the liver is reconstructed with two distinctively colored vasculature systems; namely, yellow and red vessels are observed, indicating varying oxygen saturation levels. Each one was picked up from different colored vasculature in the liver and the average value for each vasculature was calculated. As a result, the mean value was calculated to be 61% for vessels with low S-factor and 82% for vessels with high S-factor within the liver (Fig. 3(b)). Fig. 3(c) exhibits an S-factor image scanned from the dorsal side in the supine position. On the dorsal surface, dendritic vascular structures are discerned within a symmetrical 5-mm-sized oval, which can be anatomically identified as the two kidneys and their internal vascular network. Similar to the liver vascular network, this kidney vascular network also displays two types of vessels, suggesting differences in oxygen saturation. In Fig. 3(d), two vessel networks presenting a low S-factor and a singular short vessel showing a high S-factor are singled out in the kidney. The vessels with a low S-factor showcased a continuous dendritic structure, while the vessels with a high S-factor were intermittent and did not exhibit a clear branching structure. The average S-factor was 68% for both networks with low S-factors and 84% for the vessels with high S-factors.

**Fig. 3.**
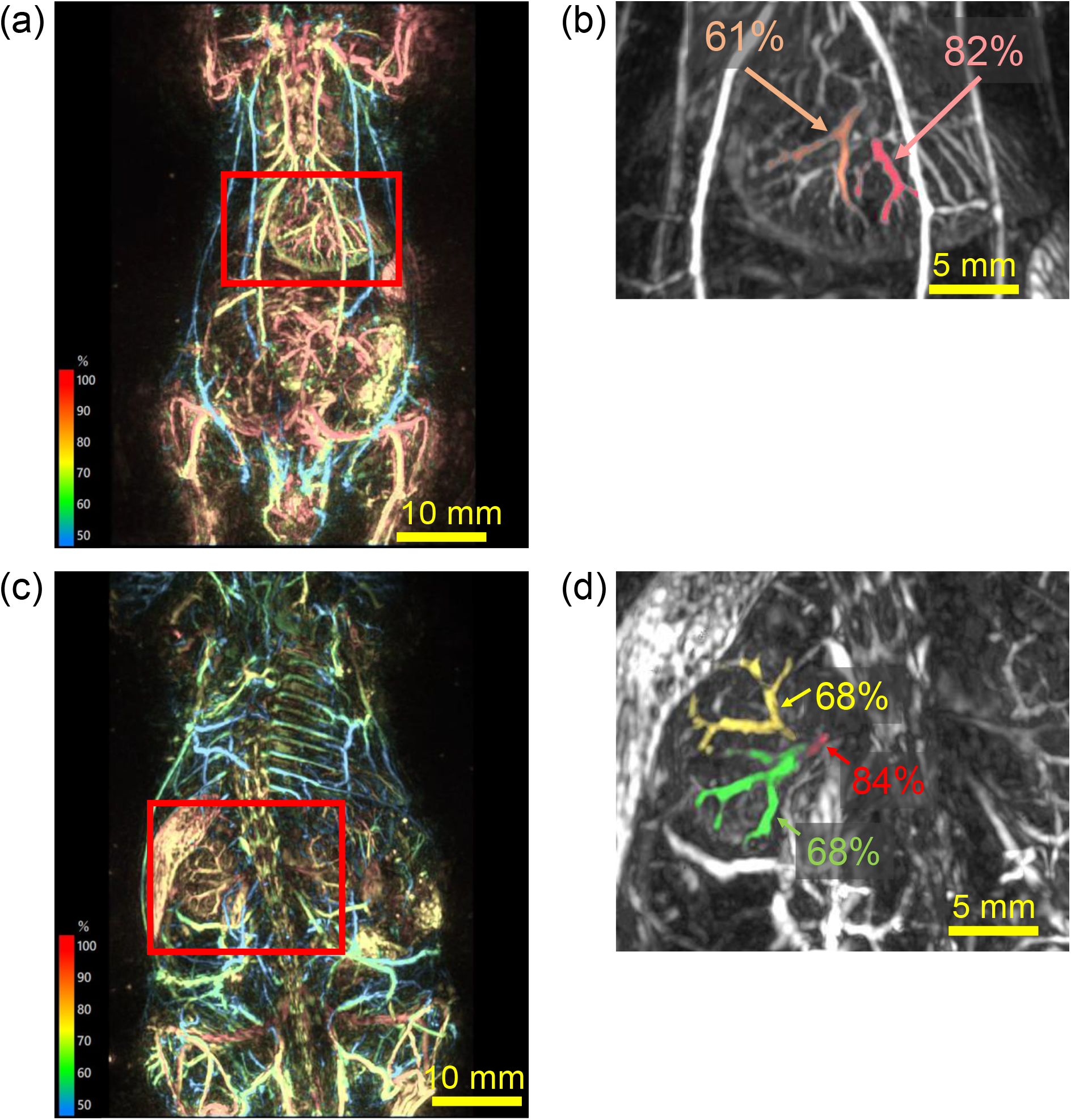
S-factor images generated by calculating photoacoustic images of mice acquired at two wavelengths, 756 nm, and 797 nm. (a) S-factor image of a mouse scanned from the ventral side within a 4 cm × 6 cm area. The red rectangle marks the liver’s location (Video-2, MP4, 10MB). (b) Average S-factor values were calculated by isolating a portion of the liver’s blood vessels denoted by the rectangle in (a). The average S-factor values for the yellow and red vessels in (a) were 61% and 82%, respectively. (c) S-factor image of a mouse scanned from the dorsal side within a 4 cm × 6 cm area. The central red rectangle marks the kidney’s location (Video-3, MP4, 7.9MB). (d) Average S-factor values were calculated by isolating a portion of the kidney’s blood vessels denoted by the rectangle in (c). The mean S-factor values for yellow and red vessels in (c) were 68% and 84%, respectively.

B6 albino mice were subsequently scanned, and image comparisons were conducted between the living and postmortem images. The resulting images are displayed in Fig. S7 and Fig. 4. Fig. S7 provides a whole-body comparison of live versus deceased B6 mice. Fig. 4 focuses on the liver and kidneys; (a)-(e) depict images of the liver, and (f)-(j) present images of the kidneys. In the liver of the in vivo condition in (a), blood vessels suggesting different oxygen saturation levels were observed, similar to the BALB/c mice in Figure 3. Vessels displaying high and low S-factor were selected to calculate the mean value of the S-factor, which was 96% and 65%, respectively, as depicted in (b). Fig. 4(c) is a post-euthanasia image of each vessel, which had transitioned to an S-factor value with no discernable arteriovenous distinction. The vessels, morphologically estimated to be identical to those in (a), had S-factor values of 41% and 36%, respectively. The S-factor values decreased significantly following death, and the differences between vessels became less pronounced. Fig. 4(e) is a graph created to visualize these respective changes.

**Fig. 4.**
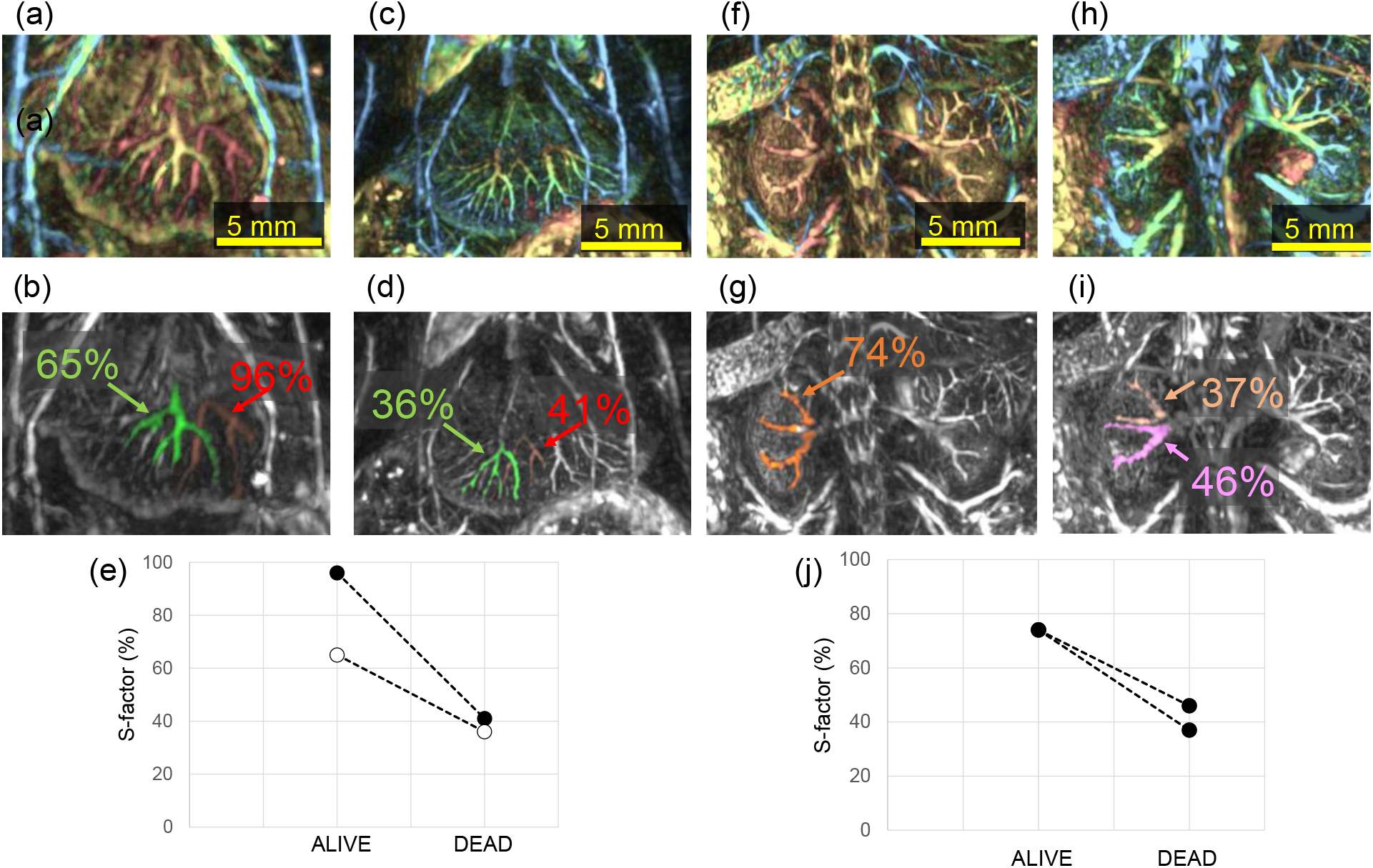
S-factor images of mice during life and post-euthanasia. (a) S-factor image of a living mouse liver, displaying variously colored vessels. (b) Average S-factor value calculated by isolating red and yellow vessels in (a). The average S-factor values for red and yellow vessels were calculated to be 96% and 65%, respectively. (c) S-factor image of the liver from the same mouse following euthanasia, as in (a) and (b). No difference in S-factor owing to vascular variation as seen in (a). (d) Average S-factor values obtained by isolating the same presumed vessels as in (b), calculated to be 41% and 36%, respectively. (e) Graphical representation of the results of (b) and (d), indicating a significant decrease in S-factor post-euthanasia. (f) S-factor image of a living mouse kidney; unlike Fig. 3(c), no differently colored vessels were observed in the kidney. (g) The average S-factor value was calculated by isolating the vessels in the kidney in (f), with the result being 74%. (h) S-factor images of the kidneys from the same mouse as in (f) and (g) post-euthanasia. Minor S-factor variations were observed depending on the vascular system. (i) Average S-factors obtained by isolating the same presumed vasculature as in (g). 37% and 46%, respectively. (j) Graphical representation of the results of (g)(i), indicating a significant decrease in S-factor post-euthanasia.

Life-death comparisons of S-factor images of the kidney region are exhibited in (f)-(j). The S-factor generally decreased after euthanasia. The color of the S-factor images was predominantly uniform, with an average value of 74%, as shown in (g). However, in the postmortem state, some vessels displayed slight differences in the color of the S-factor images as shown in (i), with mean values of 37% and 46% respectively. These values dropped to around 40%, as observed in the case of the liver.

### 3.3 Orthotopic breast cancer model mice

Orthotopic breast cancer model mice were produced by transplanting 4T1 cells into BALB/c mice. Photoacoustic images were captured daily from Day-0, the day of transplantation, to Day-11, the 11th day post-transplantation, with the exception of Day-6 due to a non-working day.

The tumor diameter increased steadily (Fig. S8), and blood vessels began to emerge in the area where the tumor cells were transplanted around the fourth day (Fig. S9). Fig. 5 illustrates an example from mouse ID #1 on day-11. Fig. 5(a) is a digital camera image captured during the scan. The tumor’s center, short diameter, and long diameter were visually identified and marked with a red circle, the diameter of which is the short diameter. This was repeated for both the left and right tumors. Fig. 5(b) depicts an S-factor image with a red circle at the same location as in (a). An orange rectangle was added to the tumor area to indicate the trimming area. Fig. 5(c) and (d) display MIP images of this area viewed from the caudal side after 3D trimming. The tumor area in (c) and (d) is a typical example, characterized by a strong signal comprising large blood vessels outside the tumor and a somewhat blurred linear structure with low S-factor value in the central region. Given that the width of (c) and (d) represents the short diameter of the tumor, the large blood vessels and areas of differing S-factor values inside the tumor were smaller than the short diameter.

**Fig. 5.**
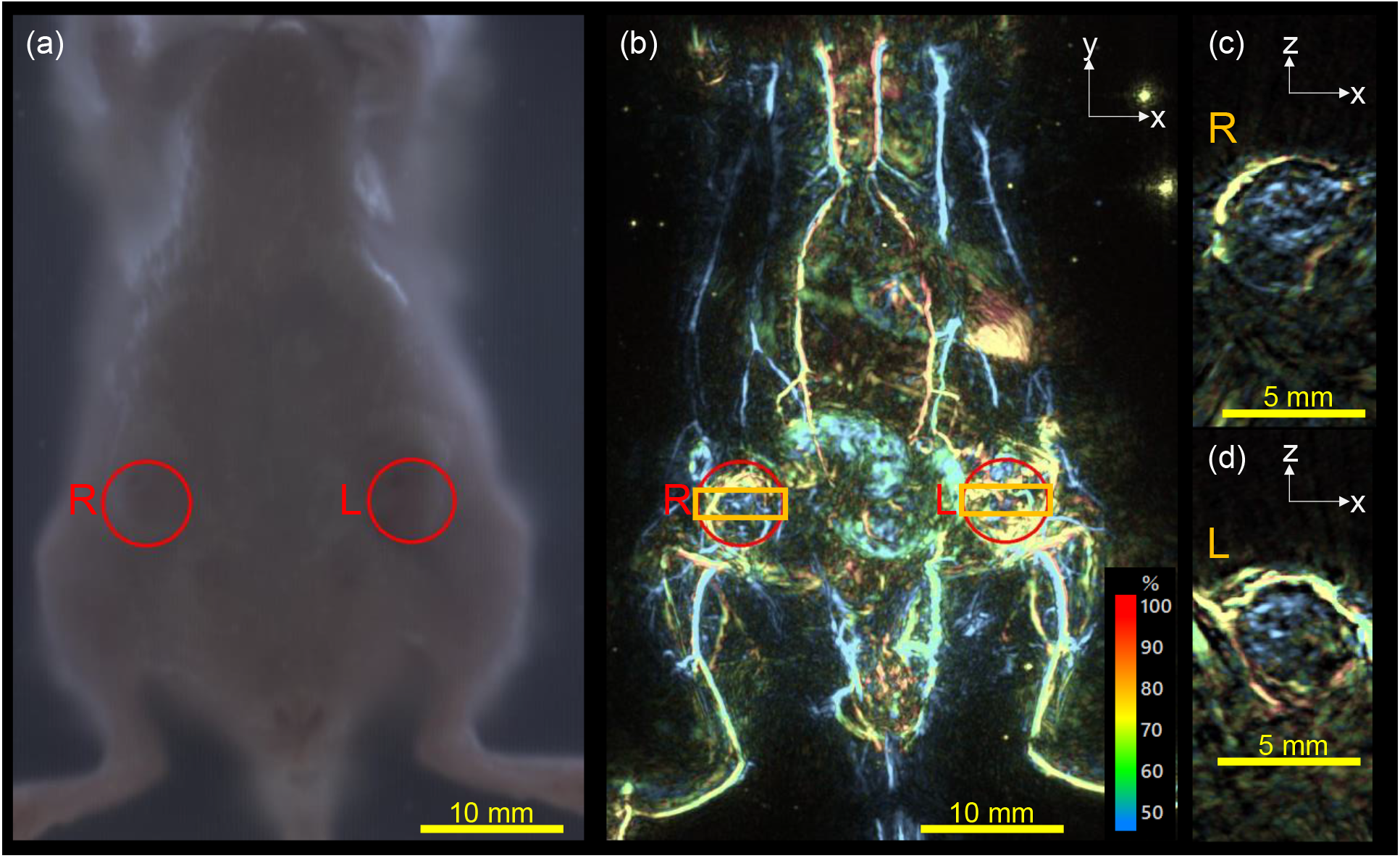
A photograph taken with a built-in digital camera and an S-factor image of an orthotopic breast cancer model mouse. (a) Photograph of a breast cancer bearing mouse taken with an integrated digital camera during scanning. The red circles indicate the right (R) and left (L) mammary gland implanted with mouse breast cancer cells. The tumor center, major diameter, and minor diameter were determined visually, and a circle with a minor diameter is shown. (b) S-factor image of (a). Red circles indicate tumors as shown in (a). The orange squares are the tumor minor diameter on the long side and 2 mm on the short side. (c) Caudal MIP image of the R mammary tumor in the orange square in (b). (d) Cross-sectional image of the left mammary tumor obtained similarly as in (c). In both (c) and (d), a thin vascular-like signal is detected inside the tumor, exhibiting a low S-factor value.

Figure 6 (MP4: 7.2MB) presents a time-sequence moving image.

**Fig. 6.**
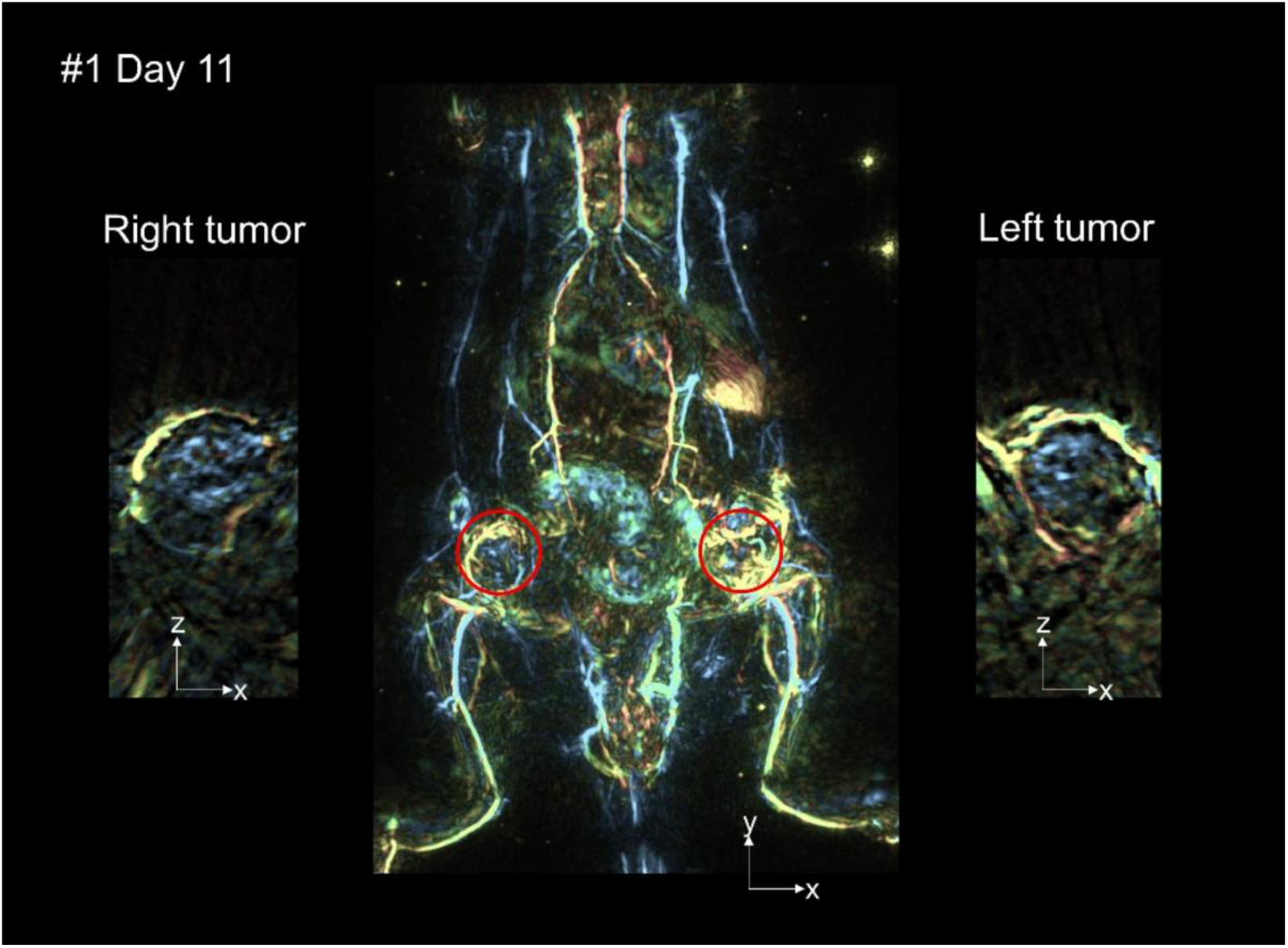
The video showcases daily changes in photoacoustic images from Day-0 to Day-11 for three mice (#1, #2, #3). The whole-body S-factor image is positioned centrally, with the tomographic image of the right breast cancer situated on the left side of the screen, and the tomographic image of the left breast cancer displayed on the right side of the screen. (Video-4, MP4, 7.2MB)

Numerical analysis of S-factor values was carried out on the right-sided tumor, as the right-sided tumor exhibited a more rounded shape than the left-sided tumor in all three mice (Fig. S9, S10). Fig. 7 reveals the results of the temporal changes in the tumor. Fig. 7(a) displays the tomogram shown in Fig. 5(c), and the yellow square outlines a rectangle used for analyzing the area inside the tumor, which is surrounded by large blood vessels in the tumor periphery. The area of the rectangle was visually determined to exclude the large vessels, and the rectangular area used for analysis was defined accordingly. Fig. 7(b) depicts the temporal changes in the average S-factor value in this region. From the image in (a), it is observable that the blue color intensifies as the tumor size increases. The graph displaying the change in S-factor value over time suggests that the S-factor value remains relatively consistent until the 8th day, before experiencing a sharp decrease after the 9th day.

**Fig. 7.**
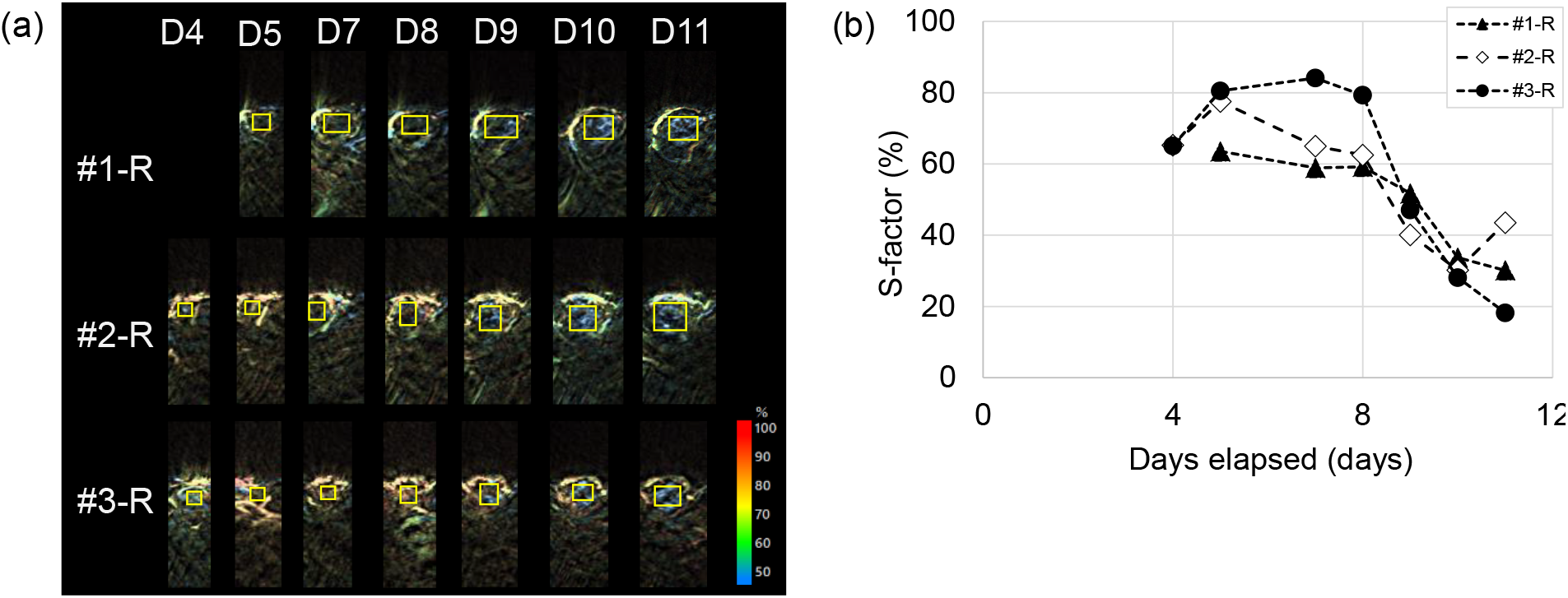
Figure showing the change in S-factor inside the tumor over time. (a) PA images showing the interior of the right mammary tumor in each of the three mice. The area inside the tumor used for analysis is indicated by yellow squares. (b) Mean values of S-factor in the area of the yellow square in (a) are plotted. The S-factor values remained nearly constant until the 8th day after transplantation of the breast cancer cell line, but a sharp decrease in S-factor values was observed from the 9th day onward.

## 4 Discussion

In this study, we demonstrated that the ISO12233 chart, a standard used widely in imaging, can effectively evaluate the performance of large-scale photoacoustic (PA) devices. This should allow for a more consistent and straightforward evaluation in line with other 2D imaging techniques, such as fluorescence imaging, which is commonly used in animal experiments. With the use of HQ mode, we were able to reduce the background noise to 1/4.6. This decrease is a factor of 16.2, considering that the number of superimpositions in the standard mode is 29 shots versus 470 shots in HQ mode. If we assume that this noise follows a white noise pattern, it’s proportional to 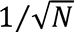 of the superimposition number, which approximates to 1/4. This 1/4.6 value for BGN is a plausible result when considering the noise as white noise.

During the mouse experiments, we employed HQ mode to obtain images with an elevated signal-to-noise ratio. We generally observed a similar morphological reproducibility of blood vessels in vivo and postmortem. This might be due to our method of lightly compressing the trunk with a weight under anesthesia, which minimizes large body movements and, consequently, reduces the influence of motion artifacts.

Two different wavelengths, 756 nm and 797 nm, were alternately irradiated, and S-factor images were subsequently reconstructed. In regards to organ blood vessels, we could observe blood vessels with varying S-factor values in any mouse strain within the liver. This suggests that arteries and veins in the liver can be distinguished. In contrast, in kidney vasculature, while we noticed a renal vasculature faintly suggesting arteries in BALB/c mice, we could not distinguish between arteries and veins in the B6 mice. This might have been because the BALB/c mice used in this study were slightly larger than the B6 mice and, therefore, had larger kidney vessels. In general, the inner diameter of arteries is smaller than that of veins. Therefore, when differentiating small arteries from veins is crucial, not only in kidney studies but also in other situations, it is recommended to use larger mice or rats, as long as the target vessels can be reconstructed.

Post-euthanasia evaluations of mice revealed a decrease in the S-factor of blood vessels throughout the body, including in the liver and kidneys. This decrease might be due to the continuation of metabolic activity in the periphery immediately after euthanasia. In this state, oxygen supply by respiration is halted due to euthanasia. For instance, it is known that patients with sleep apnea syndrome experience a decrease in oxygen saturation during sleep, and it is assumed that a similar phenomenon might occur.

The non-invasive evaluation of relative changes in S-factor over time can be repeated in the same mouse. We expect this to be applicable to various basic research and drug discovery initiatives, including the presented life-death comparisons and tumor hypoxia evaluations, as well as assessments of the effects of various drug administrations on oxygen metabolism. In particular, as this device can evaluate the S-factor of individual blood vessels, it can evaluate local oxygen metabolism in a very granular area, unlike conventional optical methods. This could potentially lead to new discoveries. However, it is vital to interpret the S-factor values presented in this study with caution, so as not to misconstrue them as absolute values of hemoglobin oxygen saturation, as noted in the text.

While morphological analysis has traditionally been the primary focus in tumor research, garnering insights from changes in oxygen saturation could provide new perspectives. Our photoacoustic imaging of the 4T1 tumor shows that, starting four days after transplantation, outer blood vessels begin to morph, forming a perimeter around the site of cancer cell implantation. Furthermore, we observed faint blood flow inside the tumor. These findings align with a similar 4T1 cell orthotopic transplantation model where CD31-positive vascular areas become identifiable on day four post-transplantation^26^. This suggests that functional blood vessels form within the tumor following a certain period after the transplantation of cancer cells.

When considering tumor hypoxia, previous research has shown that when cancer cells, transfected with a reporter gene that induces luciferase expression in the mouse body in response to the expression of hypoxia-inducible factor (HIF), are subjected to a hypoxic environment, cancer cells exhibiting high HIF activity tend to increase with tumor growth^27^. It’s important to note that this study does not solely focus on hypoxia, as HIF expression can be induced by both hypoxia and inflammatory responses. Nonetheless, our findings suggest that a more hypoxic environment might develop during specific stages of tumor growth. Pimonidazole staining is a commonly employed method to assess tumor hypoxia. Compared to tumors as small as 2 mm, pimonidazole-positive hypoxic regions have been reported to significantly expand as the tumor size increases^28^. The identification of low S-factor regions inside growing tumors seems logical, yet the sudden appearance of low S-factor regions after Day 8 warrants further investigation, including exploring the behavior of different tumor cell lines. Given the established correlation between hypoxic regions and prognosis, future developments in diagnostic methods using the S-factor could provide valuable information about highly malignant tumors, potentially informing treatment strategies.

## 5 Conclusion

We have developed a photoacoustic imaging system for animal experiments that estimates the relative oxygen saturation values for each blood vessel using two wavelengths. This system achieves distortion-free images across the entire imaging range with a spatial resolution of 0.1 mm. The S-factor images suggest the feasibility of in vivo arteriovenous vessel discrimination. By comparing images obtained before and after death, we demonstrated that oxygen saturation levels decrease postmortem. Additionally, we were able to observe temporal changes in the tumor and found that oxygen saturation decreases rapidly with increasing tumor size. Consequently, this device can be employed in a wide variety of basic research contexts.

In medical and pharmaceutical arenas, the typical research trajectory starts with basic research, proceeds to non-clinical studies using animals, and ultimately advances to clinical trials involving human subjects. As this device facilitates a seamless transition from animal experiments to clinical application in humans, it holds the potential to make significant contributions to future medical care.

## Disclosures

There is no COI.

## Code, Data, and Materials

Volume data of the vascular images shown in this paper can be provided upon request from the reader. However, the data is in Luxonus’ proprietary format and requires the dedicated software (PAT viewer) to view the 3D images.

## Acknowledgments

This work was supported in part by the ImPACT program of the Council for Science, Technology and Innovation (Cabinet Office), AMED (Grant No. 21he2302002), and NEDO (Assignment No. JPNP14012).

**Yasufumi Asao** and **Ryuichiro Hirano** contributed equally to this work.

**Yasufumi Asao** is a manager at Luxonus Inc., via a manager at Canon Inc., and an associate professor at Kyoto University, Japan. He received his BS, MS, and PhD degrees in engineering in 1991, 1993, and 2005, respectively. He is a member of SPIE.

Ryuichiro Hirano was a doctoral student at Tokyo Institute of Technology during the research period of this paper. He received his BS, MS, and PhD degrees in engineering in 2018, 2020, and 2023, respectively.

Biographies and photographs for the other authors are not available.

## Notes

### Competing Interest Statement

The authors have declared no competing interest.

### Summary of Updates

Minor correction. A space has been added between the numbers and units in the figure. Corrected the titles of the sub-items. (For example, "mice model" was corrected to "model mice" and "Photacoustic imaging of" was added.)

## References

1. J. C. Walsh, et al,. “The Clinical Importance of Assessing Tumor Hypoxia: Relationship of Tumor Hypoxia to Prognosis and Therapeutic Opportunities”, Antioxid Redox Signal., 21(10):1516–54 (2014).

2. T. Miyazaki et al., “Extracellular vesicle-mediated EBAG9 transfer from cancer cells to tumor microenvironment promotes immune escape and tumor progression”, Oncogenesis, 7(1):10 (2018).

3. I. Telarovic et al., “Interfering with Tumor Hypoxia for Radiotherapy Optimization”, J Exp Clin Cancer Res. 40: 197 (2021).

4. K. Sakata et al., “Hypoxia-induced drug resistance: comparison to P-glycoprotein-associated drug resistance.”, Br J Cancer, 64(5): 809–814, (1991).

5. Z. Fu et al., “Tumour Hypoxia-Mediated Immunosuppression: Mechanisms and Therapeutic Approaches to Improve Cancer Immunotherapy”, Cells, 1006 (2021).

6. B. Muz et al., “The role of hypoxia in cancer progression, angiogenesis, metastasis, and resistance to therapy”, Hypoxia (Auckl), (2015).

7. S. S. Cao and R. J. Kaufman, “Endoplasmic Reticulum Stress and Oxidative Stress in Cell Fate Decision and Human Disease”, Antioxid Redox Signal, 21(3): 396–413 (2014).

8. G. J. Hutchison et al., “Hypoxia-Inducible Factor 1α Expression as an Intrinsic Marker of Hypoxia: Correlation with Tumor Oxygen, Pimonidazole Measurements, and Outcome in Locally Advanced Carcinoma of the Cervix”, Clin Cancer Res., 10 (24): 8405–8412 (2004).

9. H. B. Ragnum et al., “The tumour hypoxia marker pimonidazole reflects a transcriptional programme associated with aggressive prostate cancer”, Br J Cancer. 112(2): 382–390 (2015).

10. S. Ueda et al., “Baseline Tumor Oxygen Saturation Correlates with a Pathologic Complete Response in Breast Cancer Patients Undergoing Neoadjuvant Chemotherapy”, Cancer Res 72 (17): 4318–4328 (2012).

11. Wang, L. V. and Hu, S., “Photoacoustic tomography: in vivo imaging from organelles to organs”. Science, 335(6075), 1458–62 (2012).

12. Y. Zhou et al., “Tutorial on photoacoustic tomography”, Journal of Biomedical Optics 21(6), 061007 (2016).

13. X. L. Deán-Ben et al., “Advanced optoacoustic methods for multiscale imaging of in vivo dynamics”, Chem. Soc. Rev., 46, 2158–2198 (2017).

14. Y. Xu et al., “Reconstructions in limited-view thermoacoustic tomography.”, Med Phys., 31(4), 724–33 (2004).

15. R. A. Kruger et al., “Dedicated 3D photoacoustic breast imaging.”, Med Phys., 40(11), 113301 (2013).

16. K. Nagae et al., “Real-time 3D photoacoustic visualization system with a wide field of view for imaging human limbs.”, F1000Research,. 7,1813 (2019).

17. Y. Asao et al., “In Vivo Label-Free Observation of Tumor-Related Blood Vessels in Small Animals Using a Newly Designed Photoacoustic 3D Imaging System.”, Ultrasonic Imaging, 44, 2–3 (2022).

18. Y. Asao, et al., “High-resolution photoacoustic 3D imaging system for animal experiments using a hemispherical detector array”, Proc. of SPIE Vol. 12379 123790C-9 (2023).

19. Y. Matsumoto et al., “Visualising peripheral arterioles and venules through high-resolution and large-area photoacoustic imaging.”, Sci. Rep., 8, 14930 (2018).

20. I Tsuge et al., “Preoperative vascular mapping for anterolateral thigh flap surgeries: A clinical trial of photoacoustic tomography imaging.”, Microsurgery., 40, 3, 324–330 (2019).

21. Y. Suzuki et al., “Subcutaneous Lymphatic Vessels in the Lower Extremities: Comparison between Photoacoustic Lymphangiography and Near-Infrared Fluorescence Lymphangiography.”, Radiology, 295, 469–474 (2020).

22. H. Sekiguchi and K. Togashi, “Development of the Rapid MIP Viewer for PAT data - KURUMI: Kyoto University Rapid and Universal MIP Imager.”, IEICE Tech Report, Med Imaging., 116, 163 (2017).

23. ISO 12233:2017(E), Photography — Electronic still picture imaging — Resolution and spatial frequency responses, (2017).

24. T. Kuchimaru et al. “A reliable murine model of bone metastasis by injecting cancer cells through caudal arteries”, Nat. Commun., 9, 2981 (2018)

25. V. Mehta et al., “Obstructive sleep apnea and oxygen therapy: a systematic review of the literature and meta-analysis”, J Clin Sleep Med., 9(3):271–9, (2013).

26. R. Hirano et al., “Tissue-resident macrophages are major tumor-associated macrophage resources, contributing to early TNBC development, recurrence, and metastases”, Commun Biol., 6(1):144. (2023).

27. N. T. H. Hoang et al., “Hypoxia-inducible factor-targeting prodrug TOP3combined with gemcitabine or TS-1 improves pancreatic cancer survival in an orthotopic model”, Cancer Sci., 107, 1151–1158, (2016).

28. M. Zhang et al., “Induction of Peroxiredoxin 1 by Hypoxia Regulates Heme Oxygenase-1 via NF-κB in Oral Cancer”, PLoSONE, 9(8):e105994, (2014).

