## SupplementaryFigures for "Visualization of spatial distribution of hemoglobin with various oxygen saturations in small animals using a photoacoustic imaging scanner with a hemispherical detector array"

**Fig. S1**

Schematic diagram of the newly designed experimental apparatus for animals. (a) Illustration of the bed-type apparatus previously reported, where the primary components are arranged on a flat surface, and the subject lies on the top board. (b) Illustration of the novel experimental apparatus for animals. The bed structure is bisected longitudinally, and the sensor scanning part is placed on top of the electrical components, thus forming a two-tier structure.

**Fig. S2**

Actual experimental apparatus used in this animal study. An experimenter of 1.7 m height can stand in front of the apparatus and conduct the experiment in a natural standing position.

**Fig. S3**

External view of the experimental apparatus used in this animal study. A light-shielding box is situated above the sensor to prevent the laser beam from escaping during scanning. This provision enables the device to be classified as a Class 1 laser device, removing the necessity to wear safety glasses during the experiment. Additionally, there are no location restrictions for installation.

**Fig. S4**

Spacer for the mouthpiece of an inhalation anesthesia machine used during mouse scanning. While imaging, the mouse is immersed in warm water, but the spacer prevents the mouthpiece from submerging, ensuring maintenance of breathing.

**Fig. S5**

External image of ISO 12233 test chart utilized as a phantom, inkjet printed on an A4-sized acrylic plate. The image is water-resistant, does not dissolve when immersed in water, and is printed at 1200 dpi, which is an adequate resolution for this experiment.

**Fig. S6**

Image of a mouse scanned from the abdominal side. (a) Photograph taken by the built-in digital camera, (b) Photoacoustic image captured at a single wavelength of 797 nm, and (c) Superimposed image of both (a) and (b). The combination with the digital camera image allows for clear identification of the location of blood vessels on the body.

**Fig. S7**

Comparison of images of mice both in live and post-euthanasia conditions. Both images are 4 cm wide and 6 cm high and are whole-body S-factor images. (a) Image of a live mouse scanned from the abdominal side, (b) Image of a mouse scanned post-euthanasia of the mouse shown in (a), (c) Same mouse as in (a) and (b), an image of a live mouse scanned from the back side after acquiring the image in (a), (d) Image from the back side post-euthanasia.

**Fig. S8**

Measurement of tumor diameters in three orthotopic breast cancer model mice. (a)-(c) correspond to each mouse. The long and short diameters of the left and right tumors are plotted. Tumor diameter increased in a roughly linear fashion, with no nonlinear changes observed.

**Fig. S9**

Listing of S-factor images of three orthotopic breast cancer model mice. Day 6 was a non-working day, so no experiment was conducted. It can be seen that the images are reproducible even after repeated imaging.

**Fig. S10**

List of tomographic images of the tumor area of the three orthotopic breast cancer model mice. The results of daily imaging are shown. Day 6 was a non-working day, so no experiment was conducted. The images show that blood vessels have begun to form around the tumor since around Day 4. From Day 4 to Day 8, there were blood vessels around the tumor, but the signal at the center of the tumor was thin and minute. From Day 9 onward, the blue color of the center of the tumor becomes clearer.

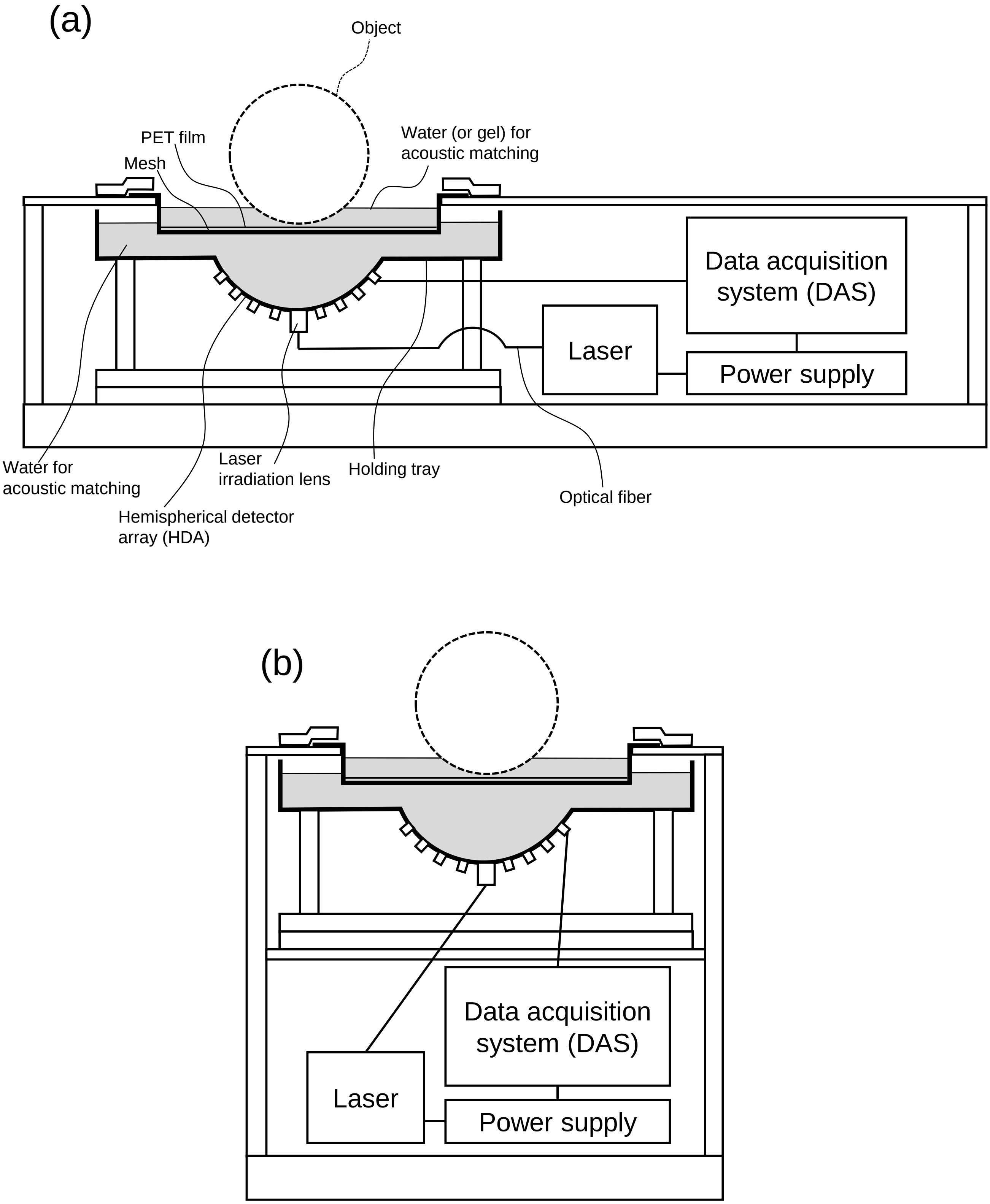
(a)
Object
PET film
Mesh
Water (or gel) for
acoustic matching
Data acquisition
system (DAS)
Laser
Power supply
Water for
acoustic matching
Laser
irradiation lens
Holding tray
Optical fiber
Hemispherical detector
array (HDA)
(b)
Data acquisition
system (DAS)
Laser
Power supply

Figure S1

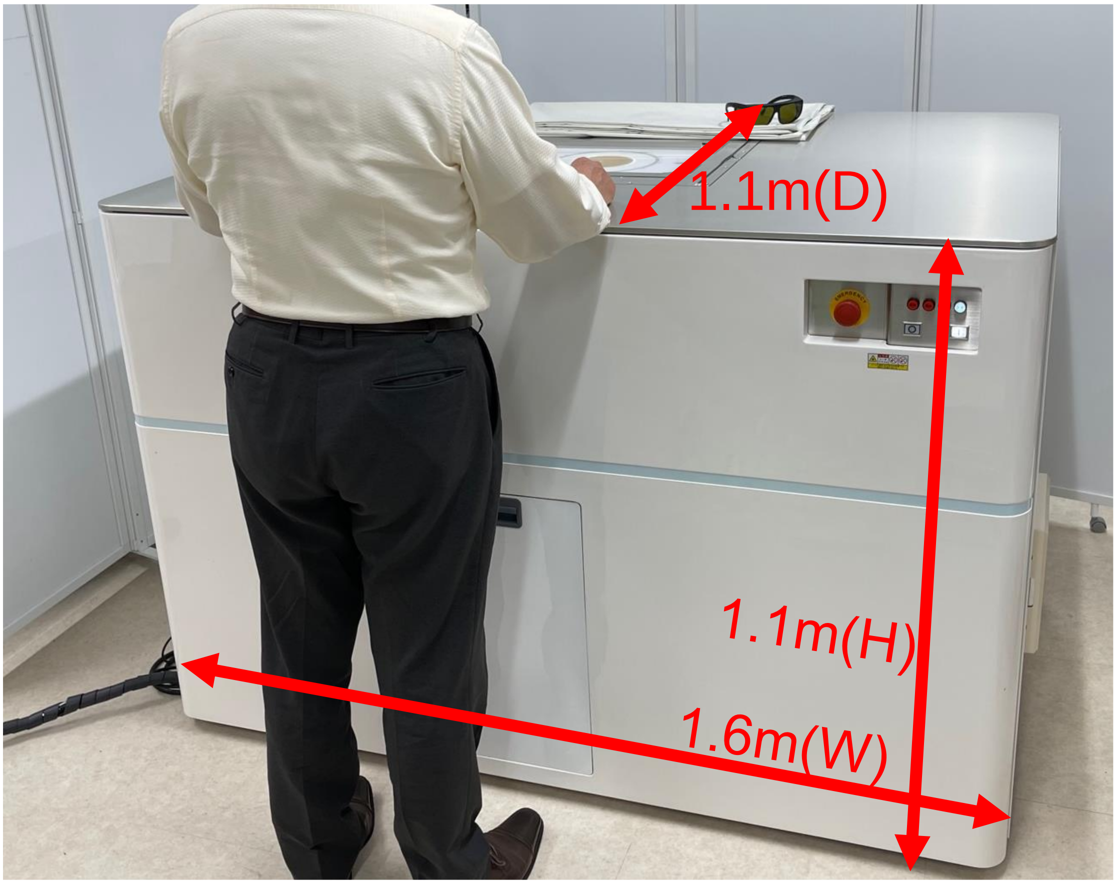


Figure S2

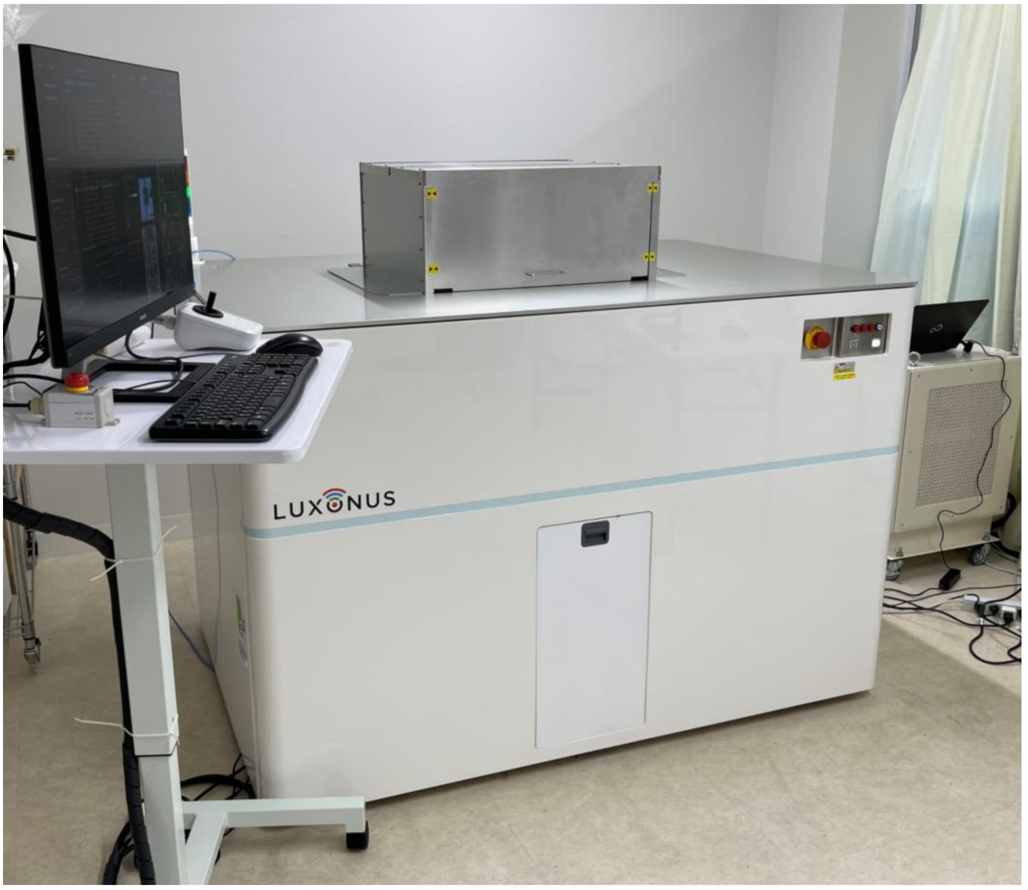


Figure S3

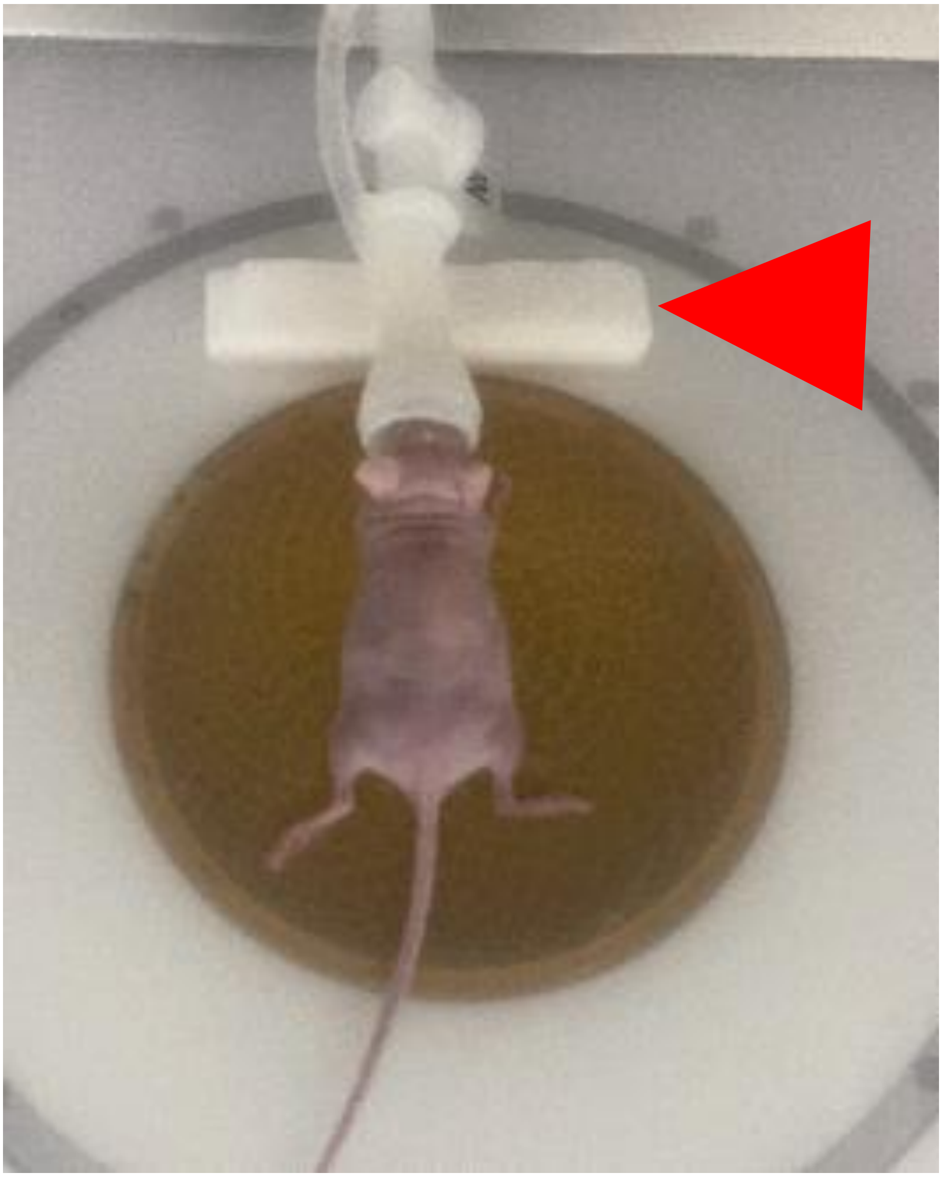

Figure S4

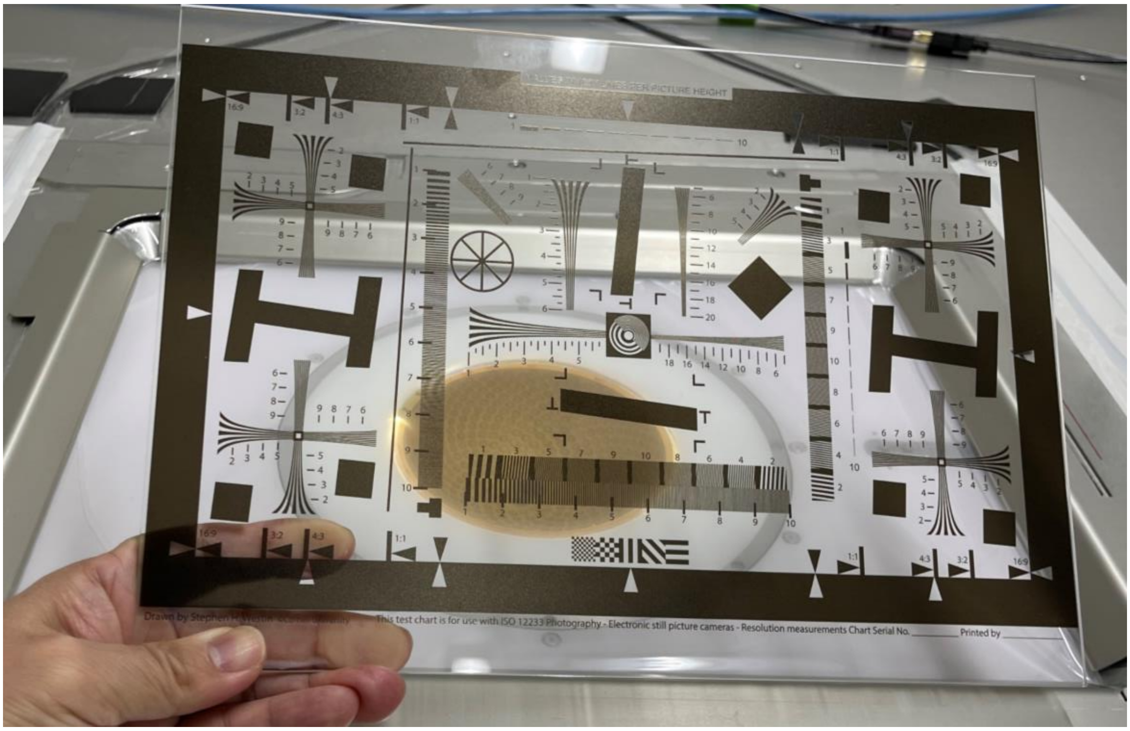
16:9
3:2
4:3
1:1
This test chart is for use with ISO 12233 Photography - Electronic still picture cameras - Resolution measurements Chart Serial No. ________ Printed by ________

Figure S5

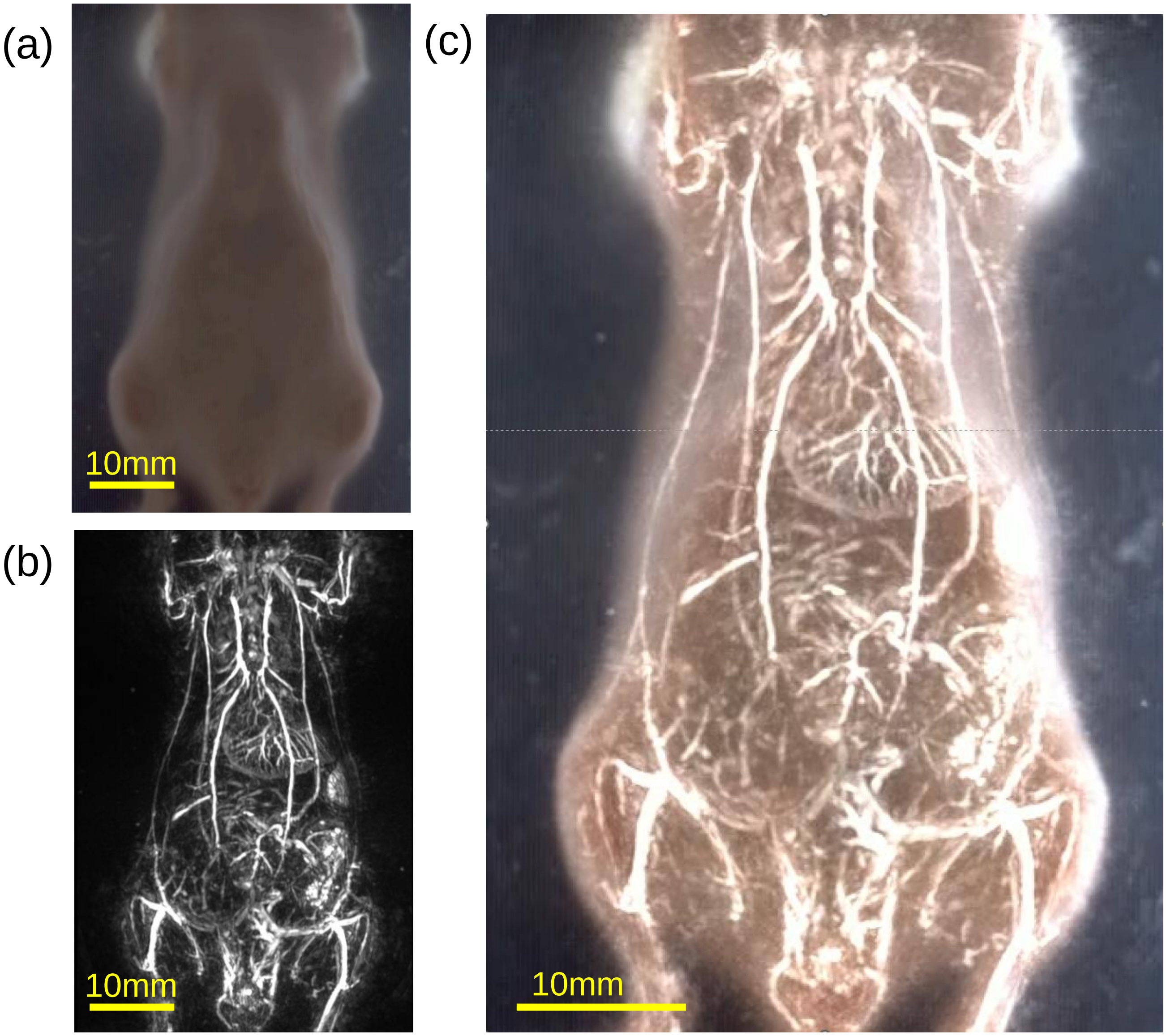


Figure S6

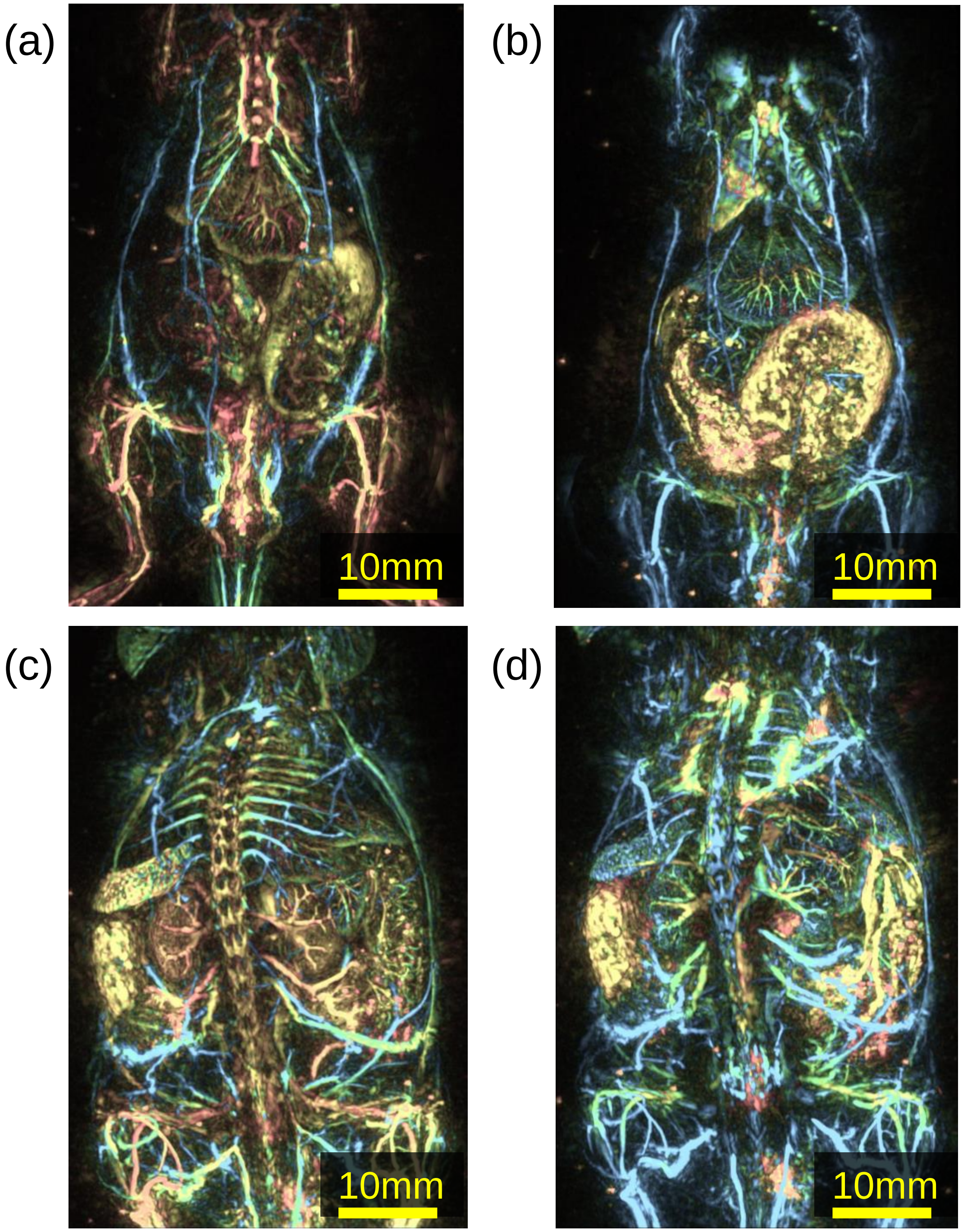


Figure S7

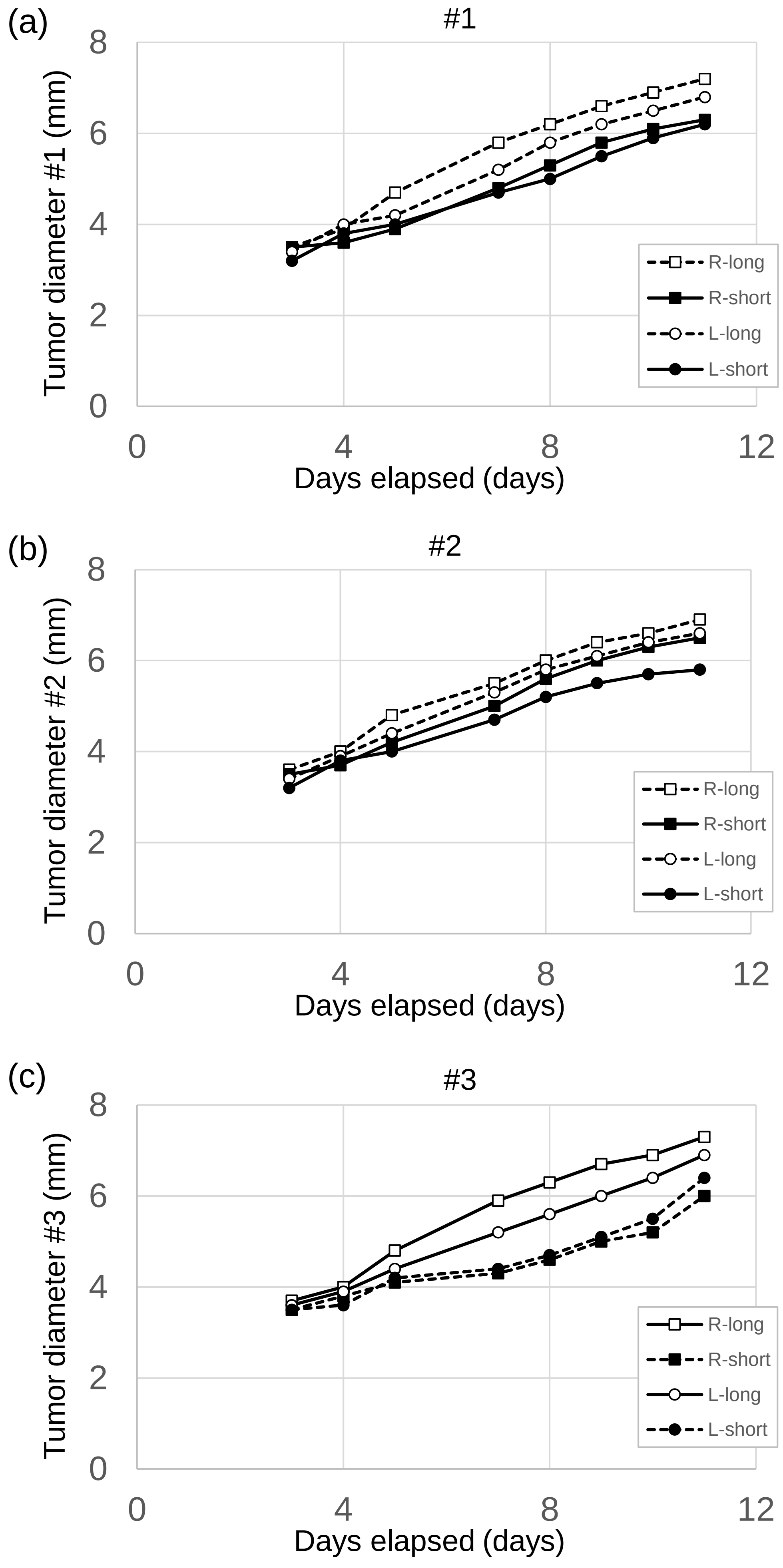
(a)
#1
Tumor diameter #1 (mm)
0
2
4
6
8
0
4
8
12
Days elapsed (days)
R-long
R-short
L-long
L-short
(b)
#2
Tumor diameter #2 (mm)
0
2
4
6
8
0
4
8
12
Days elapsed (days)
R-long
R-short
L-long
L-short
(c)
#3
Tumor diameter #3 (mm)
0
2
4
6
8
0
4
8
12
Days elapsed (days)
R-long
R-short
L-long
L-short

Figure S8

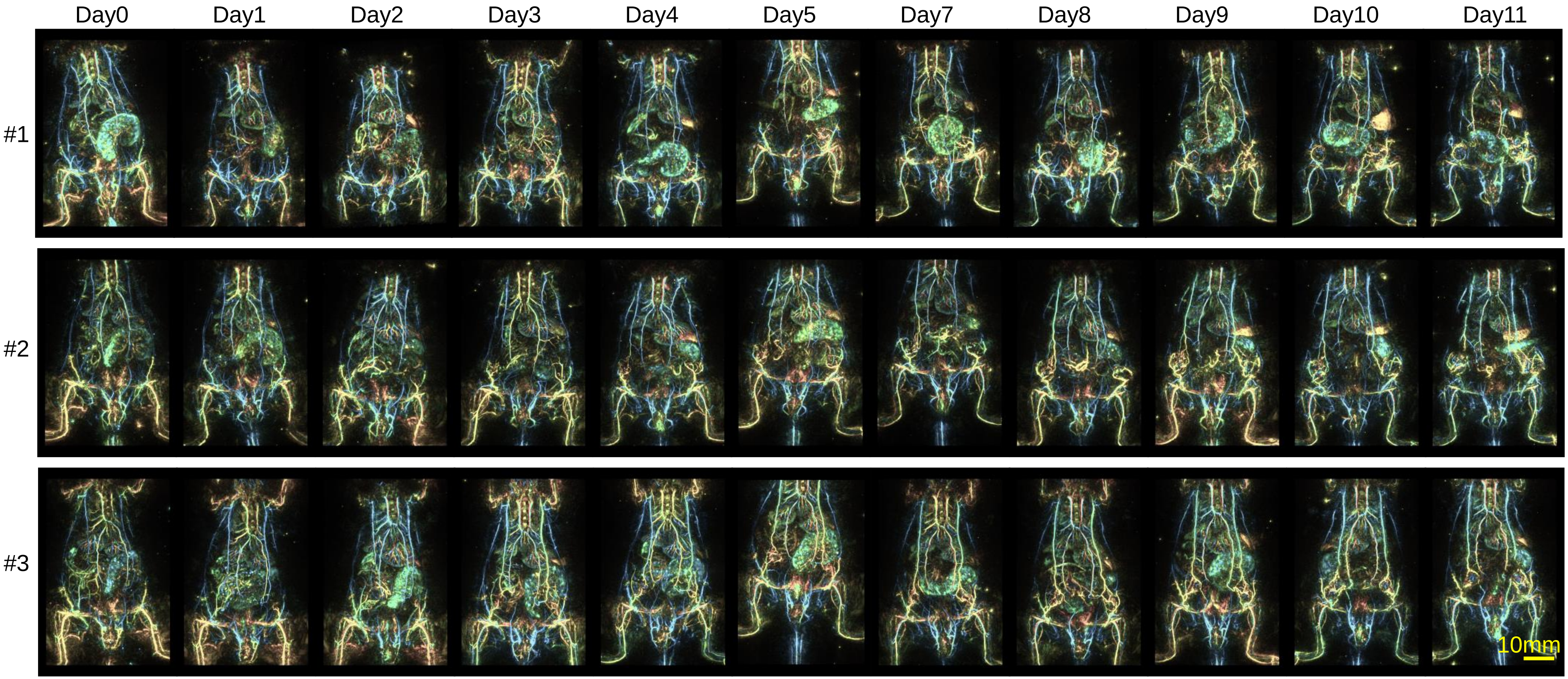
Day0
Day1
Day2
Day3
Day4
Day5
Day7
Day8
Day9
Day10
Day11
#1
#2
#3
10mm

Figure S9

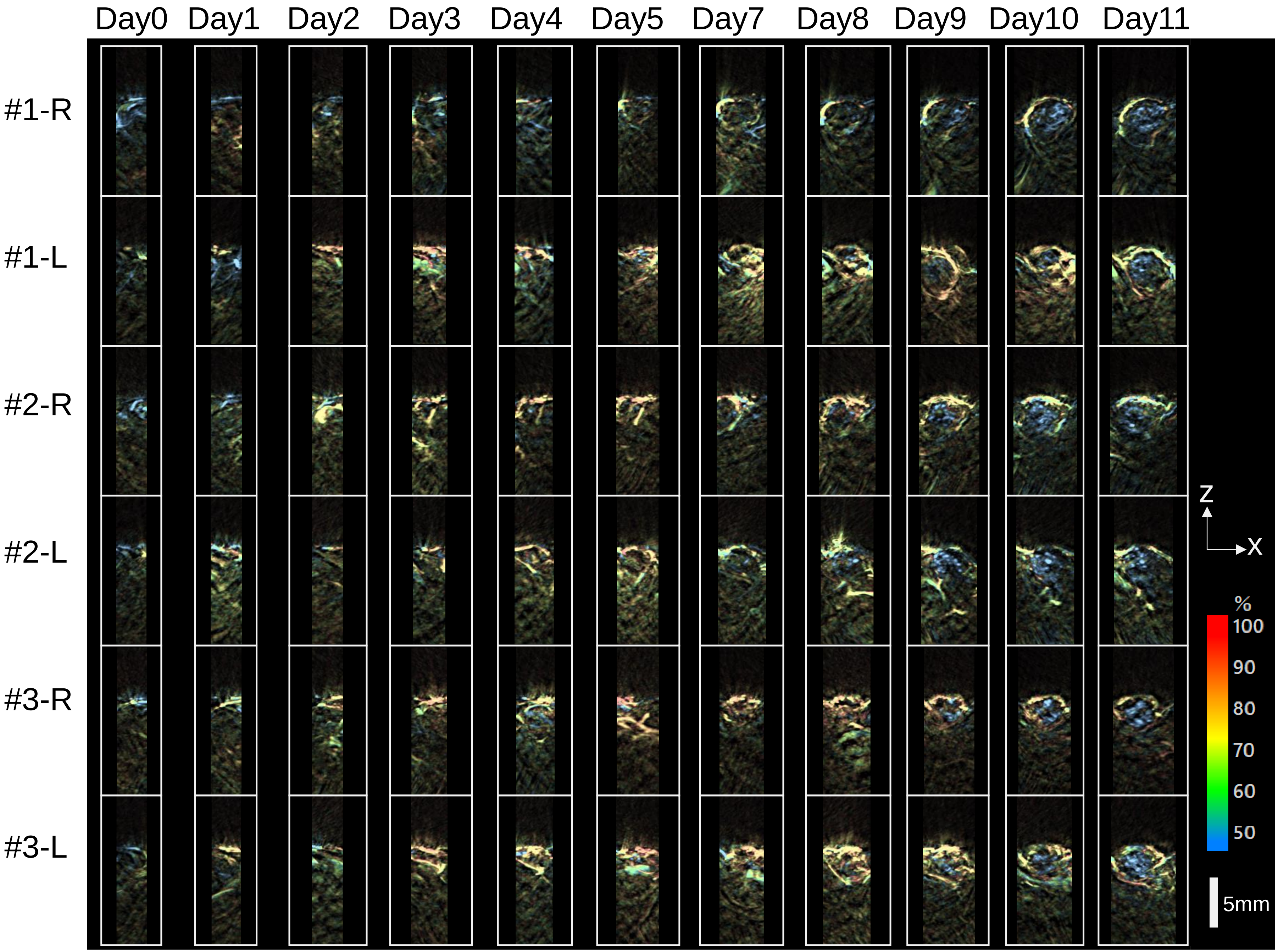
Day0
Day1
Day2
Day3
Day4
Day5
Day7
Day8
Day9
Day10
Day11
#1-R
#1-L
#2-R
#2-L
#3-R
#3-L
Z
X
%
100
90
80
70
60
50
5mm

Figure S10
